# Parallel, independent reversions to an embryonic expression phenotype in multiple types of cancer

**DOI:** 10.1101/056523

**Authors:** Corey M. Hudson, Gavin C. Conant

## Abstract

Changes in gene expression provide a valuable frame of reference for explaining the development and progression of cancer. Many tissue types radically alter their gene expression profile after becoming oncogenic. We evaluate this change in gene expression in 8 different cancer lines by comparing their expression profiles to that of their associated differentiated tissues as well as profiles for proliferative human embryonic stem cells. We find that, for non-proliferative tissues, the alterations in expression after oncogenesis result in a profile that is significantly more similar to the embryonic expression profile than to the original tissue profile. We also find that the lists of co-similar genes among embryonic and tumor cells are clustered within gene regulatory, protein interaction and metabolic networks. There is however little overlap in these lists between cancer lines and no pattern shared among all cancers in this analysis. We conclude that the manner in which cancers instantiate a proliferative pattern of expression following oncogenesis is diverse and we find no uniform proliferative program among the cancers in this analysis.

## Background

Multicellular organisms maintain numerous systems for controlling the organization and development of their constituent cells [15]. These checks are necessary in organisms that use cell differentiation to build complex organ systems and morphologies [24].Individual cells are programmed to first follow a developmental course and then assume particular functions through a combination of genomic control, epigenetic imprinting and various fate-determining signaling pathways [38]. As a result, relatively few cells in an adult multicellular organism are programmed to grow and divide without restriction [6]. However, one or a series of mutations, gene deletions, gene duplications, or epigenetic changes can break this delicate control system, resulting in proliferative cancer cells that follow a program of unrestricted division [44]. In the early stages of this change, it is expected that tumor cells have not evolved a new proliferative cellular program *ad hoc*, but through a series of mutations, primarily in the signaling and regulatory pathways, that return these cell lines to an existing proliferative program already encoded in the genome, a program that exists to facilitate embryogenesis.

However, while this general picture is reasonable, understanding the precise details by which one or more mutations give rise to the known cancer phenotypes (the genotype to phenotype mapping problem) has proven to be a distinct challenge.

Moreover, such knowledge would be of more than academic interest, as improving our understanding of this process could facilitate predictive phenotyping from tumor resequencing or improved drug design and targeting [33] [1] [41].

One approach to the problem has been genetic: the identification of risk alleles for cancer in populations. For instance, GWAS studies should help identify loci involved in the original oncogenic transition because individuals with pre-existing variation here would be at higher risk of certain cancers. However, despite their promise, the risk increase effect sizes in GWAS studies for cancers are low, with very few regions co-occuring across cancers [13]. Furthermore, studies of genomic breakpoints (i.e., common rearrangements) in resequenced cancer genomes are highly diverse and display non-overlapping patterns among cancers [28]. The determination of a specific set of genes, that in high or low copy number generally lead to oncogenesis is a current ‘dark area’ in the data from the massive cancer genome projects [14].

Strikingly, while the genetics of cancers have proven complex and dissimilar across cancer types [41], there have been observed some important common phenotypes [22]. One of the most important of these common changes is tumor cells’ switch in their primary mode of sugar metabolism. In particular, while most (resting) cells in the body prefer to respire sugars to carbon dioxide and water using oxidative phosphorylation in the mitochondria, tumor cells are much more likely to ferment those sugars using only glycolysis. This change is not minor: oxidative phosphorylation as a primary mode of metabolism appears to have been ubiquitous in the 1-2 billion year historical span covering eukaryotes and may well be the causal explanation for their uniquely complex genomes [27]. The precise importance of this *Warburg* effect is still imperfectly understood [29], but one surprising connection it suggests is to cells in the body that are *supposed* to divide rapidly:embryonic stems. These cells also display Warburg-like phenotypes [25] [37]

The extent to which this intimate connection between the metabolism of cancer and embryonic cells is the result of an epiphenomenal coincidence or a necessary functional convergence driven by natural selection pressure is unknown [2] Several studies have drawn conclusions about this relationship through the comparison of a limited number of cancers to normal tissues, but, to our knowledge, none has directly made the requisite three-way comparison of tumor, tissue and embryonic cells, despite the existence of a surfeit of next-generation sequencing and gene expression data now available.

Here, we seek to evaluate the expression profiles of various cancers with the expression profile of embryonic stem cells and adopt an explicitly network-based approach. Our goal is to evaluate the hypothesis that many tumor cells undergo a reversion to an embryonic pattern of gene expression. In principle, such a change might result from parallel changes in the expression in particular genes or by convergence at a higher organizational level.

## Methods

### Microarray Data Collection

We used gene expression data from 3 cell classes in this analysis 1) human stem cell expression data, 2) human tissue expression data and 3) associated tumor expression data. Expression data were collected from Affymetrix microarrays. To standardize the analysis, only experiments on the HG-U133̲Plus̲2 [NCBI: GPL570] platform were used Gene expression for proliferative stem cells involved 7 human embryonic stem cell lines 8 human induced pluripotent cell lines, and 2 fibroblast cell lines [NCBI GSE23402 reported in [17]]. To minimize cross-lab experimental error, only studies that included expression data from both a tumor and its associated host tissue were selected were selected. This resulted in 8 distinct cancer types (gastrointestinal cancer GSE13911, oral squamous cell carcinoma GSE30784, pancreatic cancer GSE16515,prostate cancer GSE17951, colorectal cancer GSE23878, leukemia GSE15061, breast cancer GSE10780 and lung cancer GSE19198). Each experiment had a sizeable number of independent replicates from different individuals (16-134 individuals produced the normal tissue samples and tumorous tissues were drawn from 35-181 different individuals). Affymetrix microarray experiments are prone to particular kinds of visualization errors (i.e., smears). Because of this, we manually inspected each experimental CEL file to discount the presence of smears and smudges using the affy package in Bioconductor [20]. Code for image-generation is at https://github.com/coreymhudson/AffyDistanceinthe function createImage in AffyDistance.

### Statistical comparison of expression profile distance

Each microarray experiment was normalized and error corrected using a robust multi-array average [21]. To allow values to be comparable among arrays the value for each spot intensity was then transformed by taking the intensity of *spot_i_* anddividing it by the sum of the intensity of all spots in that experiment:

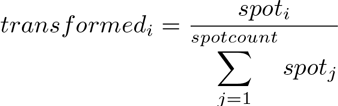
code for this transformation is in the function transformAffyData in AffyDistance.

For each probe id in each class of experiment (tumor, normal and proliferative), a 3-way pairwise comparison was made using a Kolmorogov distancemeasure (see (Algorithm 1).

**Algorithm 1:**
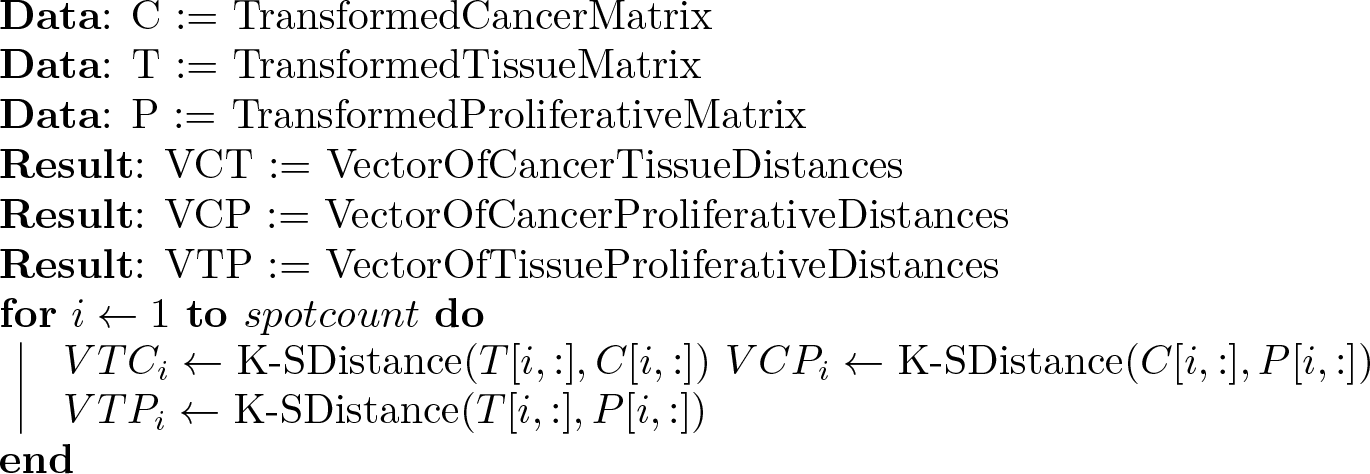
3-WayDistance

Kolmorogov distance was used because it has statistical properties that do not assume the underlying distribution is known in advance. For each cancer type (gastric, oral, pancreatic, prostate, breast, lung, leukemia and colorectal), 3 distances have been produced: cancer-normal, cancer-proliferative and normal-proliferative for each of the 54,675 probe ids in the Affymetrix HG-U133̲Plus̲2 microarray platform. Code for this transformation is built-in to the function getDistance in AffyDistance.

### P-values for lists of co-similar genes

We would like to know if the lists of co-similar embryonic and tumor genes are higher than would be expected. One null hypothesis here is that there are no genes that are significantly closer in expression between embryonic and tumor cells, when compared to both tumor and normal, (i.e., *VTC_i_*=*min(VTC_i_, VCP_i_, VTP_i_)).* A second null hypothesis is that 1/3 ofthe genes are closer in expression between embryonic and tumor cells, i.e.,: 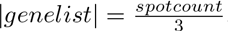. These two null hypotheses cover a continuum, going from a state in which no gene shares an expression profile between all three conditions to a state where one gene is always more similar in one of the conditions. Ideally, the co-similar genelist would contain all the genes that share an embryonic and tumor expression profile, without any spurious genes. One of the challenges in framing this test in terms of the previous null hypotheses and in comparing distances among experimental classes of different size and an unknown underlying distribution is in choosing p-values for significance in the difference in distancesαIn the presenceof multiple tests, the least conservative approach is to setαto 0.05or 0.01. Given the number of statistical tests (k=54,675), per dataset, there is a high likelihood of generating false-positives. One way of to reduce the number of potential false-positives is the Bonferroni-correction where 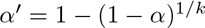 the value of which is exceedingly low for this set of experiments, of the order 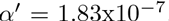. There is a high likelihood of generating false-negatives under this strategy. To minimize the trade-off between missing coexpressed genes and spuriously reported coexpression among embryonic and tumor expression profiles, we randomly reassigned the three cell classes (cancer, normal, and proliferative) for each expression value for each gene 1000 times. For each dataset, we used a 2-sample Wilcoxon-test of difference to compare the randomly reassigned”embryonic” and”cancerous” cell classes to the”normal” class. We then sought to determine the highest α-value that resulted in no pair of randomly reassigned genes being judged as statistically significant (see Figure 1). This α-value was then used for each cancer-normal paired dataset. In cases in which the cancer and embryonic expression values were found to be closer than the cancer and normal expression sets and normal and embryonic expression sets, a 2-sample Wilcoxon-test (using the previously determined α-value for significance) was used to compare the embryonic and tumor expression with the normal expression values. The genes that significantly differ in distribution were then assumed to be co-similar (see Algorithm 2 Supplemental Figure 1). Code for BootStrapGeneList is at AffyDistance in the function significantSpot.

**Algorithm 2:**
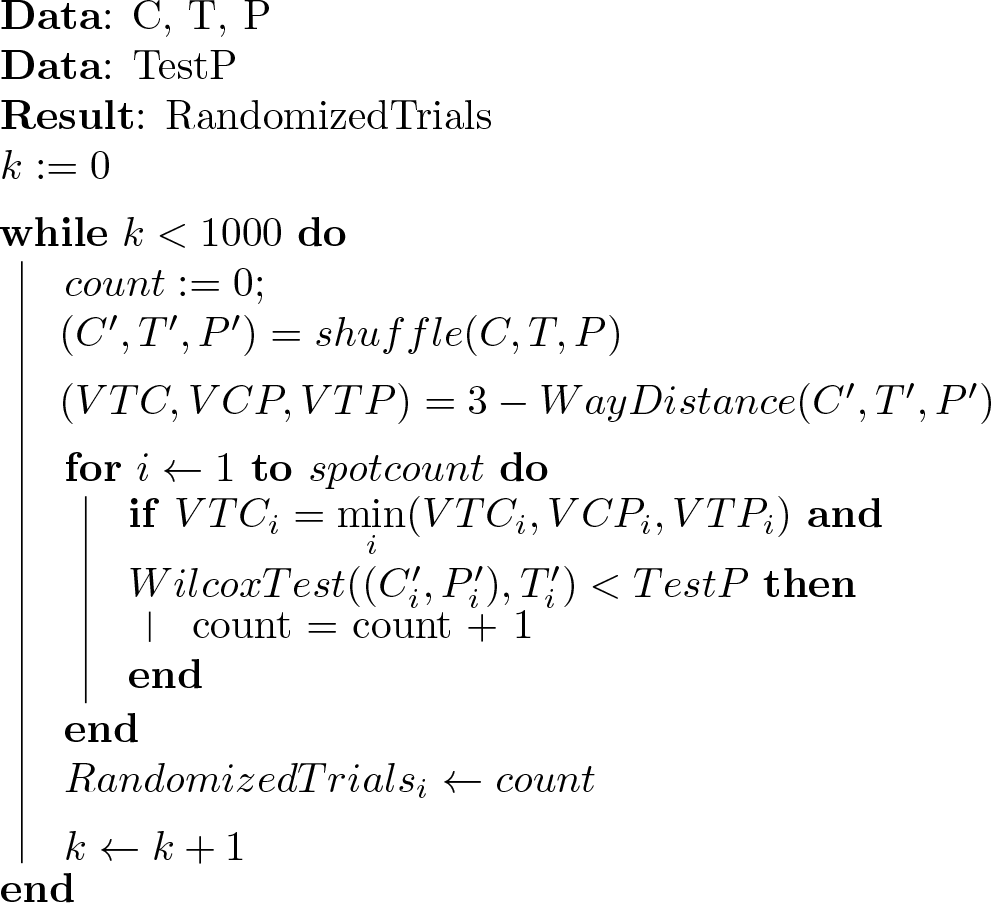
BootStrapGeneList

**Figure 1.**
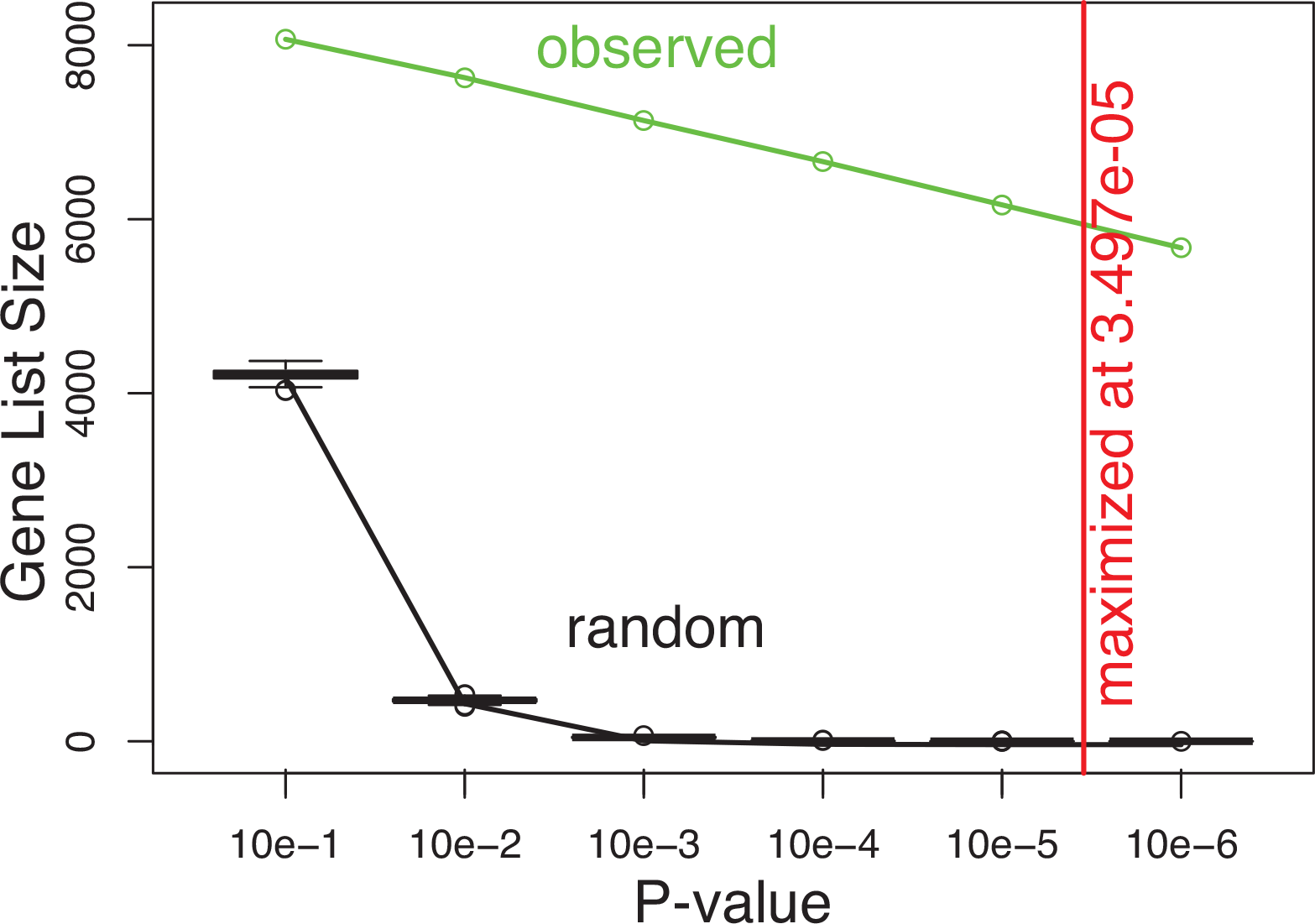
P-value minimization for 2-sample Wilcoxon-test. Embryonic and tumor vs. normal gastrointestinal tissue. The x-axis corresponds to the various p-values chosen for this analysis, the y-axis corresponds to the number of probes in the sample. The black line and boxplots illustrate a 1000 sample reshuffling of embryonic, normal and cancer classes. The green line shows the observed gene list size at given p-values. A log-log linear regression (*r*^2^=0.999, P<10e-7) of the random bootstrapped samples shows the number of probes by random error (the false-positive count) is expected to be<1 at P<3.497e-05. At this p-value the observed number of probes is 6351.

### Network evaluation

We used four networks in our evaluation of tumor/embryonic co-expression (protein-protein [34], gene regulatory [39], metabolic [11], and functional annotation [19]). The goal of our network analysis is to ascertain if the shared genes of a pair of cell classes also cluster in these networks [12]. Since protein-interactions, metabolic reactions and gene regulation all work in concert to form the cells underlying machinery [23], we also evaluated the combination of the protein interaction (PPI), metabolic (MN) and regulatory networks(GRN). This *combined network* (hereafter CN) is formally defined formally as *G(v, e)*=⋃*edges*⊃{*PPI, MN, GRN*}.

We used several methods to evaluate the clustering in these networks. We measured the transitivity (also known as the average clustering coefficient [43]) for the CN, PPI, MN, and GRNs. We also measured the number of connected components [18]. The statistical significance of these values were evaluated by bootstrapping 107 random iterations of the network and recalculating these statistics. Fully random networks tend to be a poor representation of real-world networks [7]. One of the primary characteristics of real networks are their power-law degree distributions [46]. Our randomization preserved the number of interactions for each node, while randomized which nodes interacted. This allowed us to retain each networks power-law degree distributions while still randomizing the topology [42].

In addition to measuring transitivity and the number and size of connected components (all of which can be measured directly), we also evaluated the fit of these networks into highly interconnected communities [31]. The methods for detecting these are not exact, since the fit of vertices into communities is known to be NP-hard [5]. The strength of communities was evaluated using a modularity statistic, which essentially measures the number of edges within communities versus the number of edges between communities There were several classes of heuristics used in this approximation of maximum modularity [26]. Since weare essentially choosing among heuristics we implemented several of these classes, including *iterative removal of edges based on betweenness* [32], *greedy modularity maximization* [8], *label propagation* [36], and *random walk* [35] methods. The statistical significance of these was evaluated by generating 107 randomized degree-preserving networks and calculating the maximum modularity for using each heuristic for both the observed and random networks. Network analysis was conducted using the igraph package in R [9]

### Functional analysis of network neighbors

We took the list of coexpressed genes for the combined network for each cancer type and evaluated the over-representation of functional classes among the largest 3 communities from the using the DAVID Bioinformatics Resource [19]. We limited the annotations to the Gene Ontology Biological Process and Metabolic Function annotations, and KEGG Pathway annotations. We ranked and evaluated the significance of annotations using Benjamini p-values (one of a variety of False Discovery Rate (FDR) minimization techniques-this version of the FDR does not assume independence of gene lists [3]), which are robust to multiple tests, false positives and hierarchical annotations and evaluated the 10 highest ranking annotation clusters [40]. The statistical significance for any given community in the network was evaluated by taking the number of edges within the community for each node and the number of edges between communities for each node and calculating a Wilcox rank sum statistic.

## Results

### Statistical comparison of expression profile distances

We found that expression distances between cancers and embryos were closer than expression distances between normal tissues and embryos for most genes in almost all the cancers (excluding pancreatic cancer: see Table 1). This being despite the fact that cancer and normal tissue expression values were collected from the same lab and embryo expression measurements were taken in numerous other labs. This trend suggests a pattern of shared expression between cancer and embryo for most genes. To evaluate the statistical significance of this trend,we used a binomial test with the null hypothesis that the tumor and normal tissue cell classes were equally likely to have genes that were close to the embryonic pattern (e.g., 50% of the time the tumor would be closer vs. 50% of the time normal tissue would be closer). For tissues that can be said to be proliferative in their healthy tissue state (white-blood cell and pancreatic B-cell) the proportion of spots where the expression distance between embryo and cancer is less than the expression distance between embryo and healthy tissue varies between 0.481 and 0.543. For cancers in which the associated healthy tissues are non-proliferative (colorectal, oral squamous, prostate, gastrointestinal, breast, and lung cancers) these proportions range from 0.568 to 0.664 and are all statistically greater than 0.5 (i.e., genes are more likely to be similar in expression between tumor and embryonic cells than normal and embryonic cells; P<0.001, Table 1).

**Table 1.** Distance measures between cancer, normal and embryonic cells among proliferative (i.e., leukemia and pancreatic) associated tissue and non-proliferative associated tissues (i.e.,colorectal, oral squamous, prostate, gastro-intestinal, breast and lung).

| Cancer type | Number of cases of tumor and embryonic expression profiles being closest | Number of cases of tissue and embryonic expression profiles being closest | Proportion (tumor/tissue) |
| --- | --- | --- | --- |
| Colorectal | 31549** | 19650 | 0.616 |
| Oral squamous | 34683** | 17946 | 0.659 |
| Prostate | 32978** | 18184 | 0.645 |
| Gastro-intestinal | 27933** | 21176 | 0.568 |
| Breast | 28418** | 18598 | 0.604 |
| Lung | 35580** | 17950 | 0.664 |
| *Leukemia* | *28665\*\** | *24152* | *0.543* |
| *Pancreatic* | *24345* | *26269* | *0.481* |

\*\*statistically significant for Binomial difference in equal proportions (proportion = 0.5) at 0.001

### Lists of co-similar genes

For each of the non-proliferative tissues, the empirically determined α-values that evaluate whether similarity in expression between two classes is statistically significant are of a similar order (from 2.07e-05to 5.77e-05). They are all also close to roughly 2 orders of magnitude higher than the Bonferroni-corrected ±^´^-values (1.83e-07). The number of genes that were found to be co-similar between the cancer cells and embryonic cells varies between 5514 and 9972 (Table 2 and Figure 1). The expected number of genes in these lists is<1 (P<0.001) and are based on 1000 random reassignments of cancer, normal and embryonic expression values. These data strongly suggests that there is a much larger than expected gene cohort in which the expression profile between cancer and embryonic cell types are more similar than between cancer and tissue cell types in the cancers of non-proliferative tissues.

**Table 2.** The number of probes with co-similar expression between tumor and embryonic tissue for each cancer type at *P-values* determined to have fewer than 1 false positive.

| Cancer type | Significant probes | Empirically determined $P$-values |
| --- | --- | --- |
| Gastrointestinal | 6351 | $3.49\text{e-}05$ |
| Oral squamous | 6394 | $2.07\text{e-}05$ |
| Colorectal | 6625 | $5.77\text{e-}05$ |
| Prostate | 8959 | $3.25\text{e-}05$ |
| Breast | 5514 | $3.27\text{e-}05$ |
| Lung | 9972 | $3.07\text{e-}05$ |

### Gene overlap

Given the similarities between six different non-proliferative cancers and the embryonic cell samples, one might expect that a common set of genes would have changed in expression across these six cancers. However, our results do not illustrate this trend: for the 6 sets of experiments reported in this study, no one gene was shared across the embryonically-similar sets of all six cancers. These 19,210 genes are enriched for an embryonic expression profile in at least 1 experiment. This includes 8,916 that are more highly expressed than in the normal tissue and 10,294 that are more lowly expressed in the normal tissue. The overlap among experiments is considerably lower. With 2465 genes sharing expression between 2 or more experiments, 285 sharing expression between 3 or more experiments, 36 sharing expression between 4 or more experiments, and 0 sharing expression in 5or more experiments. The decrease in overlap is similarly dramatic when the Affymetrix spots associated with these genes are mapped onto Uniprot proteinids. When the Uniprot enzymes and transporters are mapped to the H.sapiens Recon 1 metabolic model [11], the overlap decreases dramatically as well, with the exception that 2 reactions(K+-Cl-cotransport and 3’,5’-cyclic-nucleotide phosphodiesterase) which overlap expression in 5 different cancers (see Figure 2).

**Figure 2.**
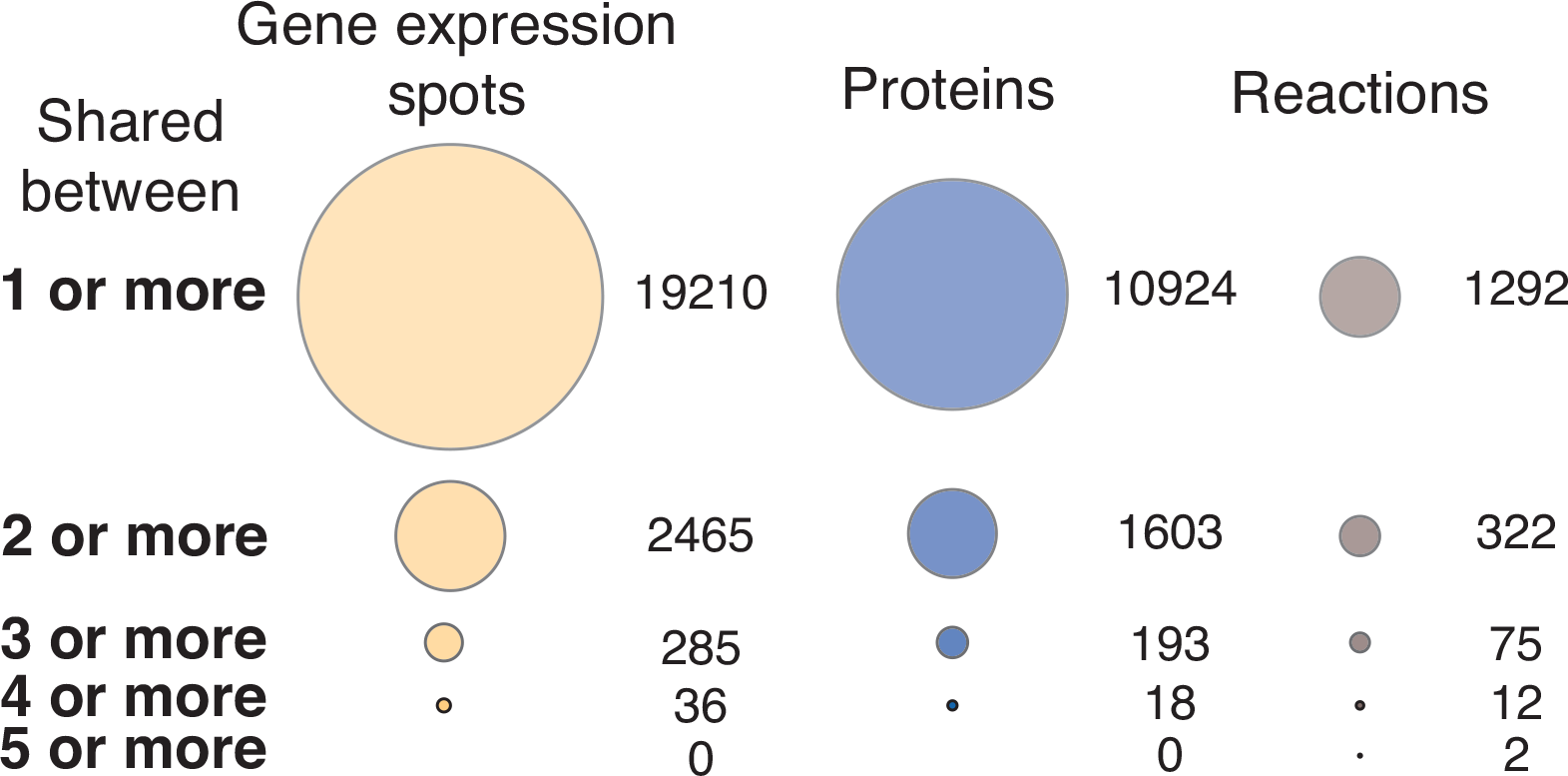
Overlap in significant gene lists among different cancer types for genes, proteins, and reactions. The size of spots corresponds tothe number of probes shared between two experiments. Gene expression spots are from Affymetrix HG-U133̲Plus̲2 Microarrays. Protein values have been mapped onto Uniprot IDs. Reaction values have been mapped onto the H. sapiens Recon 1 metabolic model.

### Cancer networks

For each of the 6 cancers in non-proliferative tissues, the combined networks, protein-interaction networks and metabolic networks have higher than expected average transitivity (P<1e-06), meaning that the co-similar gene lists in these networks form tight-knit interacting clusters (see Supplemental Table 1). All of the combined networks, protein-interaction networks and metabolic networks also have a smaller than expected number of clusters (P<0.01). This suggests a dense clustering (i.e., a small number of large, highly interacting clusters), of genes that change in expression upon conversion to an oncogenic phenotype (Figure 3).

Unlike the previous three networks, the gene regulatory networks behave very differently. In particular, the gene regulatory networks have either non-significant or lower than expected transitivity and a lower than expected number of gene clusters (P<1e-06). The source of this difference may lie in the structural differences between regulatory networks and the other types of networks considered. Thus, it appears that regulatory networks are seldom highly interconnected [16]because, unlike protein-interaction and metabolic networks that have interacting functional modules,gene regulatory networks instead show a strongly hierarchical structure in addition to being modular. For these networks modularity refers to a distinct and non-overlapping groups of co-regulated genes and their shared regulators.

**Figure 3.**
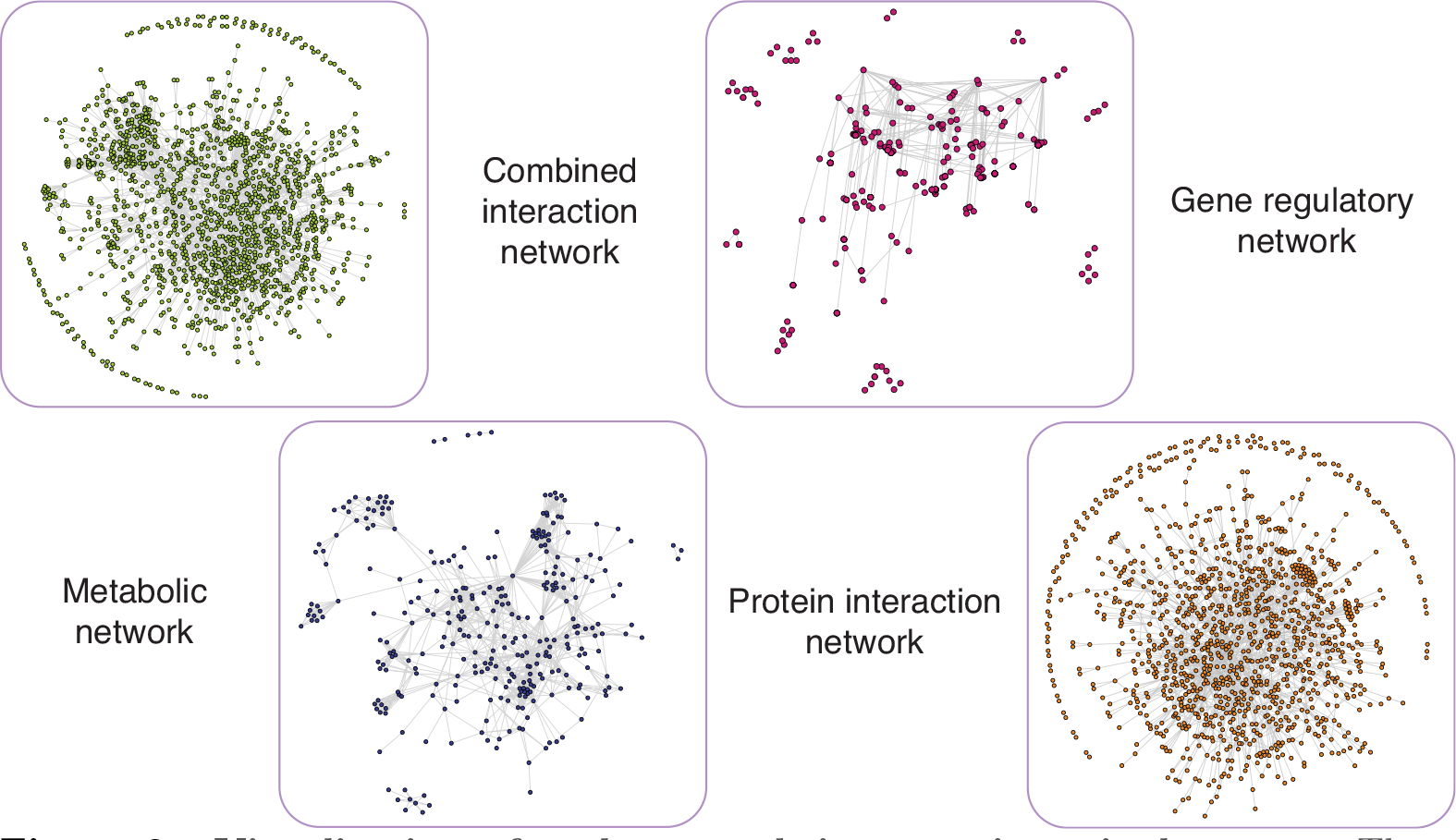
Visualization of each network in gastrointestinal cancer. These visualizations show the fundamental features of each of these four networks. The gene regulatory network is has a small number of clusters and is not very highly interconnected. The protein interaction network has one large very highly interconnected cluster and many small satellite clusters (mostly made of pairs of proteins). The metabolic network hasvery fewclusters, which are highly modular and highly interconnected. The combined network has one large highly interconnected cluster and many smallsatellite clusters.

Each of the 4 heuristics for estimating modularity (*greedy, edge betweenness, label propagation, and random walk*) strongly support the hypothesis of modularity across the 4 network types(Supplemental Table 1). In other words, each type of network, whatever their other differences in structure,tend to consist of distinct units with few interconnections between those units.

### Annotation of network features

The 3 largest combined network clusters share many of the same annotation categories across all 6 cancers in non-proliferative tissues (see Supplemental Table 2). All of the 3 largest network clusters are statistically significant (within cluster edges > between cluster edges: Wilcox Test: P<2.2e-16). Taking the 10highest scoring annotation clusters (based on Benjamini p-value), there are 44 categories shared between all 6 types (Supplemental Table 3). These fall broadly into the categories: transcription, nucleic acid metabolism, regulation of biosynthesis, andATP binding; all of which are primary cellular functions. There are also 21 categories that are shared by 5 cancer types (see SupplementalTable 4), which fall broadly into the categories: apoptosis, mitotic cell cycle, and phosphorylation. There are also115 categories unique to each cancer (see Supplemental Table 5).This includes categories like “negative regulation of DNAbinding”, which is a specialization of transcription and DNA binding; uninformative categories like “spliceosome”; and categories like “mTOR signaling” which are expected to be important in both oncogenesis and embryonic stem cell differentiation [45] [10].

## Discussion

In this study, we found that breast, colorectal, gastrointestinal, lung, oral squamous and prostate cancers showed a distinct expression pattern similar to the expression pattern in embryonic cells. This strategy gives us a window into the genetic underpinnings of proliferative behavior in cancer. We find that the genes that share expression between cancer and embryonic cells form distinct clusters. This occurs in terms of gene regulation, protein interaction, and metabolism and suggests that these clusters are functionally significant. Despite this similarity in the formation of gene clusters, the clusters themselves and the genes of similar expression underlying them show, very little overlap between the different types of cancer. It is unknown whether this lack of overlap is due to the random nature of oncogenic events, (e.g., mutation, gene duplication and deletion, or epigenetic changes) the selective microenvironment in which the cell resides or the limited overlap in expression among the original associated tissue. However, each of these cancers express a large set of genes in patterns similar to those seen in embryos Despite this similarity, we find very few patterns emerging in cancers generally. This is not solely a function of scale, since we consider variation in gene, protein, and metabolic reaction. At each of these scales, the overlap sometimes shared across two or more cancer types but rarely across more than that.

We assert that cancer cells are individuals, from an evolutionary point of view [30], and that cancer phenotypes are, at that scale, not only functional, but potentially selectively advantageous [4]. This presents something of a paradox. This year millions of people will get cancer. Yet, the manner in which cancer emerges is due to complex interactions between a large number of heterogeneous external factors (smoking, solar rays, pollutants, etc.) and various internal genetic predispositions. Importantly, the initial cancer or tumor development takes a relatively short period of time (as measured in numbers of cell divisions) and hence occurs in a small population of cells. Given this relatively limited space for evolution to operate, it may be surprising that cancers are often able to dramatically change their expression profiles and phenotypes.

One possible explanation for why cancers do rapidly evolve and share so many aspects of their phenotype (the so-called *hallmarks of cancer*)is that cancer is the result of a small and simple set of aberrant genetic/protein/metabolic changes. Our results argue against this, as do the low effect sizes among GWAS studies. We find very little overlap in gene expression among the cancers in our study, whether we consider individual gene coexpression, proteins co-occurrence, or metabolic interactions. We hope that further analysis will be able to follow up this work and evaluate the extent to which the similarities between the programs of proliferation in embryos and tumors are superficial or causal. We also hope that work in this area will lead to a more thorough and mechanistic understanding of the manner in which cancerous cells canalized to preexisting embryonic phenotypes. Such work would help in understanding whether these observations of an ‘embryonic gene program’ are phenomenological or preconditioned.

## Acknowledgments

Thanks to J. Chris Pires, Dmitry Korkin, Jailin Cheng, Dustin Mayfield and Patrick Edger for reading and commenting on early drafts of this work. Thanks also to the NLM Bioinformatics and Health Informatics Training Fellowship and the Reproductive Biology Group of the Food for the 21st century program at the University of Missouri who funded this research.

## Supplemental Figures and Tables

### Supplemental Figures

**Figure S1.**
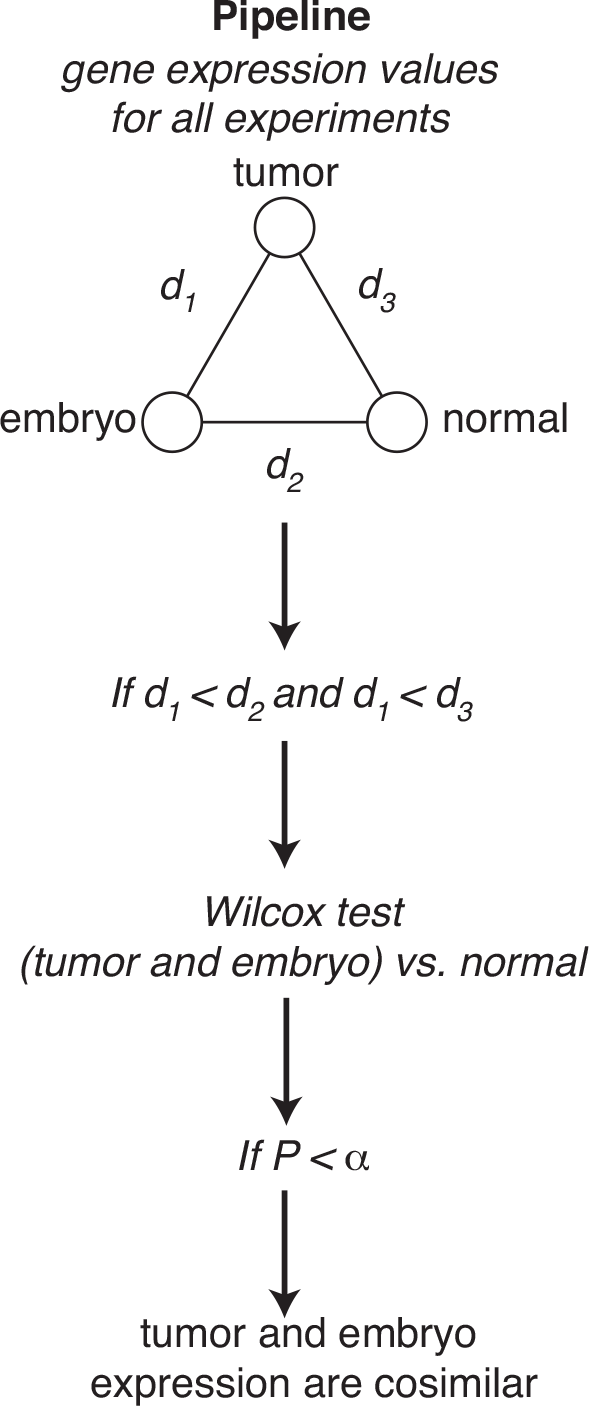
Pipeline for statistical analysis of tumor / embryo co-similarity. *d*_1_ *d*_2_, *d*_3_ correspond to Kolmorogov distance between the expression vectors for tumors, embryos and associated healthy cells. If *d*_1_ is less than *d*_2_and is less than *d*_3_ perform a Wilcox test. This test is for both embryo and cancer expression vs. normal expression. If the value is less than α (see Figure 1), the expression is co-similar between tumor and embryo.

### Supplemental Tables

**Table S1.**
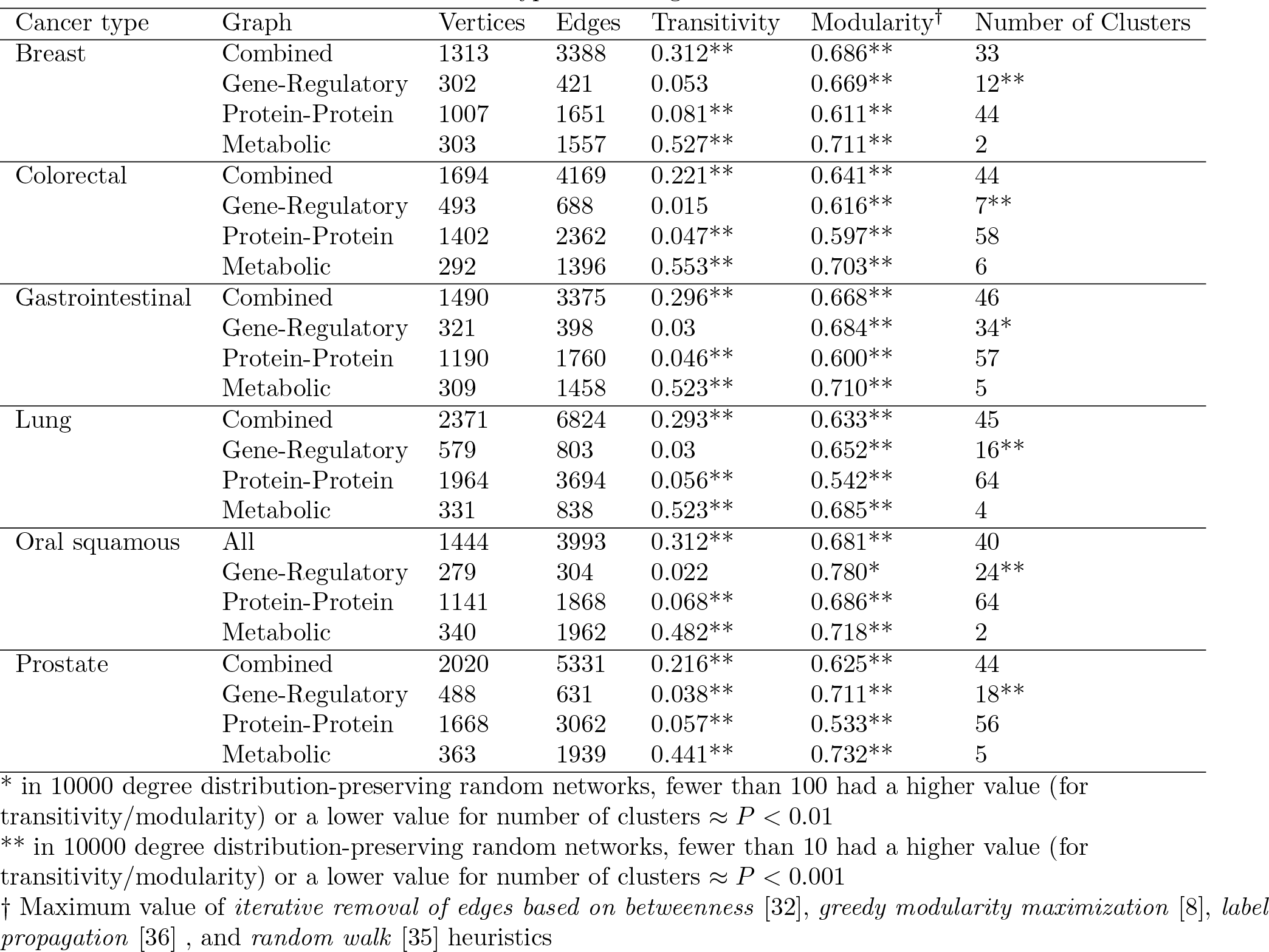
Network statistics for each cancer type and biological network

| Cancer type | Graph | Vertices | Edges | Transitivity | Modularity<sup>†</sup> | Number of Clusters |
| --- | --- | --- | --- | --- | --- | --- |
| Breast | Combined | 1313 | 3388 | 0.312** | 0.686** | 33 |
|  | Gene-Regulatory | 302 | 421 | 0.053 | 0.669** | 12** |
|  | Protein-Protein | 1007 | 1651 | 0.081** | 0.611** | 44 |
|  | Metabolic | 303 | 1557 | 0.527** | 0.711** | 2 |
| Colorectal | Combined | 1694 | 4169 | 0.221** | 0.641** | 44 |
|  | Gene-Regulatory | 493 | 688 | 0.015 | 0.616** | 7** |
|  | Protein-Protein | 1402 | 2362 | 0.047** | 0.597** | 58 |
|  | Metabolic | 292 | 1396 | 0.553** | 0.703** | 6 |
| Gastrointestinal | Combined | 1490 | 3375 | 0.296** | 0.668** | 46 |
|  | Gene-Regulatory | 321 | 398 | 0.03 | 0.684** | 34* |
|  | Protein-Protein | 1190 | 1760 | 0.046** | 0.600** | 57 |
|  | Metabolic | 309 | 1458 | 0.523** | 0.710** | 5 |
| Lung | Combined | 2371 | 6824 | 0.293** | 0.633** | 45 |
|  | Gene-Regulatory | 579 | 803 | 0.03 | 0.652** | 16** |
|  | Protein-Protein | 1964 | 3694 | 0.056** | 0.542** | 64 |
|  | Metabolic | 331 | 838 | 0.523** | 0.685** | 4 |
| Oral squamous | All | 1444 | 3993 | 0.312** | 0.681** | 40 |
|  | Gene-Regulatory | 279 | 304 | 0.022 | 0.780* | 24** |
|  | Protein-Protein | 1141 | 1868 | 0.068** | 0.686** | 64 |
|  | Metabolic | 340 | 1962 | 0.482** | 0.718** | 2 |
| Prostate | Combined | 2020 | 5331 | 0.216** | 0.625** | 44 |
|  | Gene-Regulatory | 488 | 631 | 0.038** | 0.711** | 18** |
|  | Protein-Protein | 1668 | 3062 | 0.057** | 0.533** | 56 |
|  | Metabolic | 363 | 1939 | 0.441** | 0.732** | 5 |

\* in 10000 degree distribution-preserving random networks, fewer than 100 had a higher value (for transitivity/modularity) or a lower value for number of clusters $\approx P < 0.01$
\*\* in 10000 degree distribution-preserving random networks, fewer than 10 had a higher value (for transitivity/modularity) or a lower value for number of clusters $\approx P < 0.001$
<sup>†</sup> Maximum value of *iterative removal of edges based on betweenness* [32], *greedy modularity maximization* [8], *label propagation* [36], and *random walk* [35] heuristics

**Table S2.**
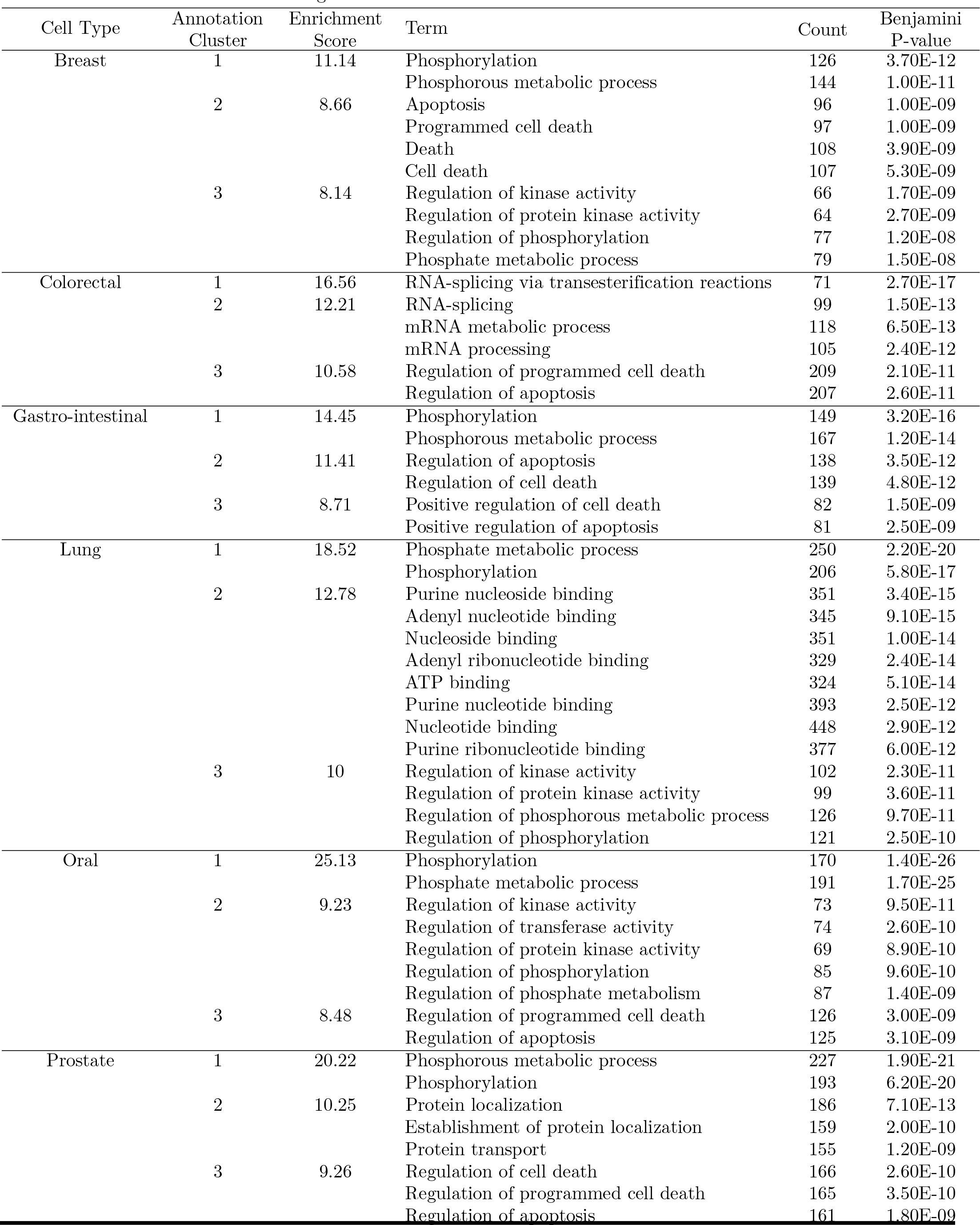
Annotation Term Clustering.

**Table S3.** Terms in All Six Cell Types

| Functional annotation | Number of shared cell types |
| --- | --- |
| GO:0016563~transcription activator activity | 6 |
| GO:0051172~negative regulation of nitrogen compound metabolic process | 6 |
| GO:0051173~positive regulation of nitrogen compound metabolic process | 6 |
| GO:0032559~adenyl ribonucleotide binding | 6 |
| GO:0016481~negative regulation of transcription | 6 |
| GO:0032555~purine ribonucleotide binding | 6 |
| GO:0003712~transcription cofactor activity | 6 |
| GO:0030554~adenyl nucleotide binding | 6 |
| GO:0051254~positive regulation of RNA metabolic process | 6 |
| GO:0010628~positive regulation of gene expression | 6 |
| GO:0032553~ribonucleotide binding | 6 |
| GO:0045449~regulation of transcription | 6 |
| GO:0006357~regulation of transcription from RNA polymerase II promoter | 6 |
| GO:0010605~negative regulation of macromolecule metabolic process | 6 |
| GO:0001883~purine nucleoside binding | 6 |
| GO:0031327~negative regulation of cellular biosynthetic process | 6 |
| GO:0010604~positive regulation of macromolecule metabolic process | 6 |
| GO:0045893~positive regulation of transcription; DNA-dependent | 6 |
| GO:0010557~positive regulation of macromolecule biosynthetic process | 6 |
| GO:0045944~positive regulation of transcription from RNA polymerase II promoter | 6 |
| GO:0003700~transcription factor activity | 6 |
| GO:0031328~positive regulation of cellular biosynthetic process | 6 |
| GO:0009891~positive regulation of biosynthetic process | 6 |
| GO:0010629~negative regulation of gene expression | 6 |
| GO:0000122~negative regulation of transcription from RNA polymerase II promoter | 6 |
| GO:0000166~nucleotide binding | 6 |
| GO:0017076~purine nucleotide binding | 6 |
| GO:0045934~negative regulation of nucleobase; nucleoside; nucleotide and nucleic acid metabolic process | 6 |
| GO:0016564~transcription repressor activity | 6 |
| GO:0001882~nucleoside binding | 6 |
| GO:0008134~transcription factor binding | 6 |
| GO:0003713~transcription coactivator activity | 6 |
| GO:0010558~negative regulation of macromolecule biosynthetic process | 6 |
| GO:0045892~negative regulation of transcription; DNA-dependent | 6 |
| GO:0005524~ATP binding | 6 |
| GO:0006355~regulation of transcription; DNA-dependent | 6 |
| GO:0006350~transcription | 6 |
| GO:0009890~negative regulation of biosynthetic process | 6 |
| GO:0045935~positive regulation of nucleobase; nucleoside; nucleotide and nucleic acid metabolic process | 6 |
| GO:0051253~negative regulation of RNA metabolic process | 6 |
| GO:0051252~regulation of RNA metabolic process | 6 |
| GO:0045941~positive regulation of transcription | 6 |
| GO:0003677~DNA binding | 6 |
| GO:0030528~transcription regulator activity | 6 |

**Table S4.** Terms in Five Cell Types

| Functional annotation | Number of shared cell types |
| --- | --- |
| GO:0006796~phosphate metabolic process | 5 |
| GO:0016265~death | 5 |
| GO:0022403~cell cycle phase | 5 |
| GO:0000278~mitotic cell cycle | 5 |
| GO:0006917~induction of apoptosis | 5 |
| GO:0010942~positive regulation of cell death | 5 |
| GO:0010941~regulation of cell death | 5 |
| GO:0008219~cell death | 5 |
| GO:0012501~programmed cell death | 5 |
| GO:0043068~positive regulation of programmed cell death | 5 |
| GO:0006468~protein amino acid phosphorylation | 5 |
| GO:0006915~apoptosis | 5 |
| GO:0042981~regulation of apoptosis | 5 |
| GO:0043067~regulation of programmed cell death | 5 |
| GO:0004672~protein kinase activity | 5 |
| GO:0016310~phosphorylation | 5 |
| GO:0012502~induction of programmed cell death | 5 |
| GO:0043065~positive regulation of apoptosis | 5 |
| GO:0007049~cell cycle | 5 |
| GO:0004674~protein serine/threonine kinase activity | 5 |
| GO:0006793~phosphorus metabolic process | 5 |
| GO:0022402~cell cycle process | 5 |

**Table S5.** Terms in Five Cell Types.

| Functional Annotation | Number of shared cell types |
| --- | --- |
| GO:0043086~negative regulation of catalytic activity | 1 |
| GO:0016573~histone acetylation | 1 |
| GO:0032990~cell part morphogenesis | 1 |
| GO:0000.95~RNA splicing; via transesterification reactions | 1 |
| GO:00.9059~chromosome segregation | 1 |
| GO:0016407~acetyltransferase activity | 1 |
| GO:0016071~mRNA metabolic process | 1 |
| hsa0.922:Neurotrophin signaling pathway | 1 |
| hsa04666:Fc gamma R-mediated phagocytosis | 1 |
| GO:0060537~muscle tissue development | 1 |
| GO:000.913~protein tyrosine kinase activity | 1 |
| GO:0033673~negative regulation of kinase activity | 1 |
| GO:0051099~positive regulation of binding | 1 |
| GO:0014706~striated muscle tissue development | 1 |
| GO:0004402~histone acetyltransferase activity | 1 |
| hsa03040:Spliceosome | 1 |
| GO:00.9411~axon guidance | 1 |
| GO:0010740~positive regulation of protein kinase cascade | 1 |
| GO:0032355~response to estradiol stimulus | 1 |
| GO:0060538~skeletal muscle organ development | 1 |
| hsa04150:mTOR signaling pathway | 1 |
| GO:0018212~peptidyl-tyrosine modification | 1 |
| GO:00.9519~skeletal muscle tissue development | 1 |
| GO:0016337~cell-cell adhesion | 1 |
| GO:0031175~neuron projection development | 1 |
| GO:0048666~neuron development | 1 |
| hsa04660:T cell receptor signaling pathway | 1 |
| GO:0048667~cell morphogenesis involved in neuron differentiation | 1 |
| GO:0043543~protein amino acid acylation | 1 |
| GO:0010647~positive regulation of cell communication | 1 |
| GO:0008080~N-acetyltransferase activity | 1 |
| GO:0030522~intracellular receptor-mediated signaling pathway | 1 |
| GO:0051092~positive regulation of NF-kappaB transcription factor activity | 1 |
| hsa04012:ErbB signaling pathway | 1 |
| GO:0043388~positive regulation of DNA binding | 1 |
| hsa04662:B cell receptor signaling pathway | 1 |
| GO:0030.94~protein binding; bridging | 1 |
| GO:0006352~transcription initiation | 1 |
| GO:00.9169~transmembrane receptor protein tyrosine kinase signaling pathway | 1 |
| hsa04650:Natural killer cell mediated cytotoxicity | 1 |
| GO:0006469~negative regulation of protein kinase activity | 1 |
| GO:0046332~SMAD binding | 1 |
| GO:00.9155~cell adhesion | 1 |
| GO:0000.97~RNA splicing; via transesterification reactions with bulged adenosine as nucleophile | 1 |
| GO:0004112~cyclic-nucleotide phosphodiesterase activity | 1 |
| GO:0030518~steroid hormone receptor signaling pathway | 1 |
| GO:0004114~3';5'-cyclic-nucleotide phosphodiesterase activity | 1 |
| GO:0051091~positive regulation of transcription factor activity | 1 |

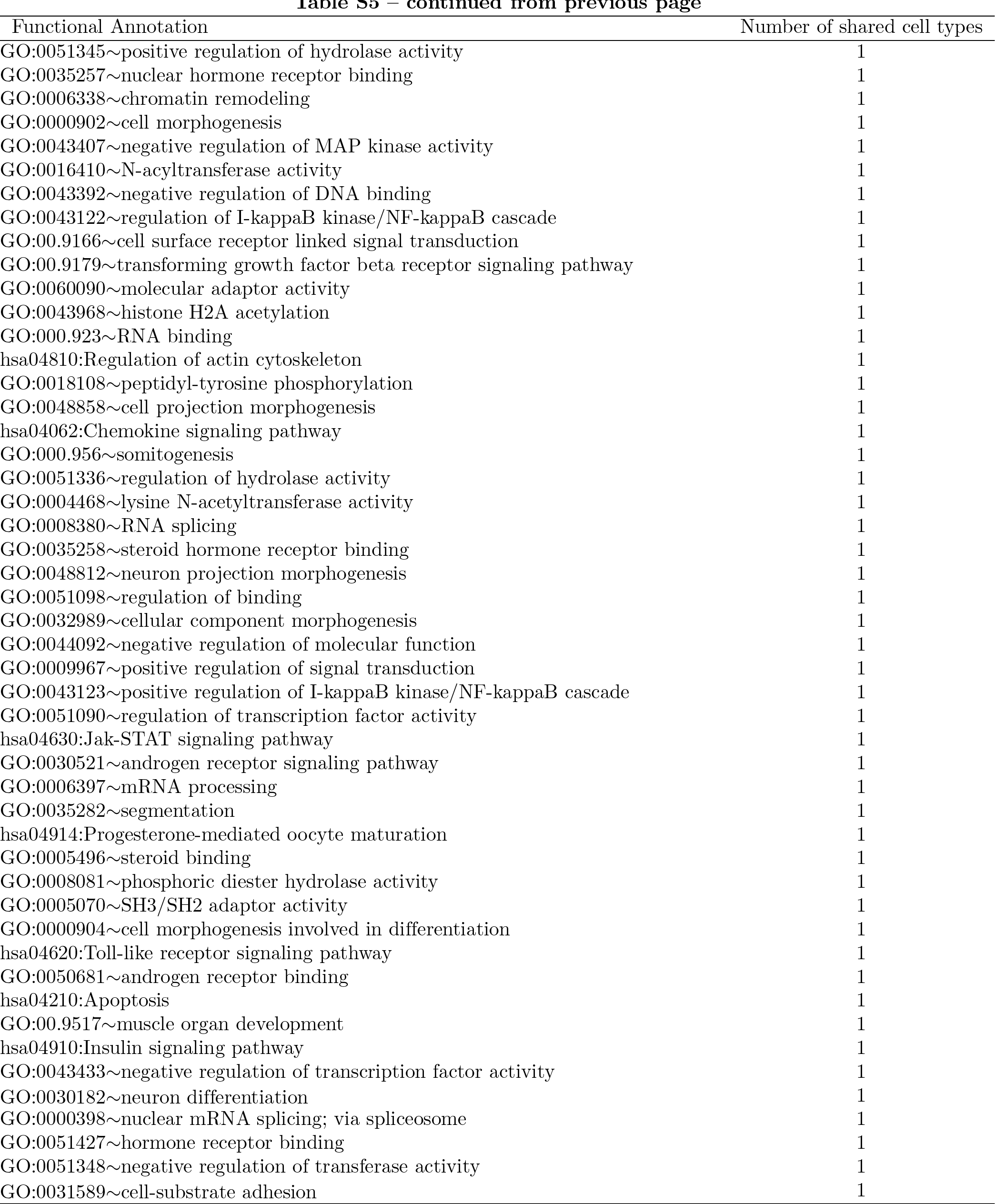

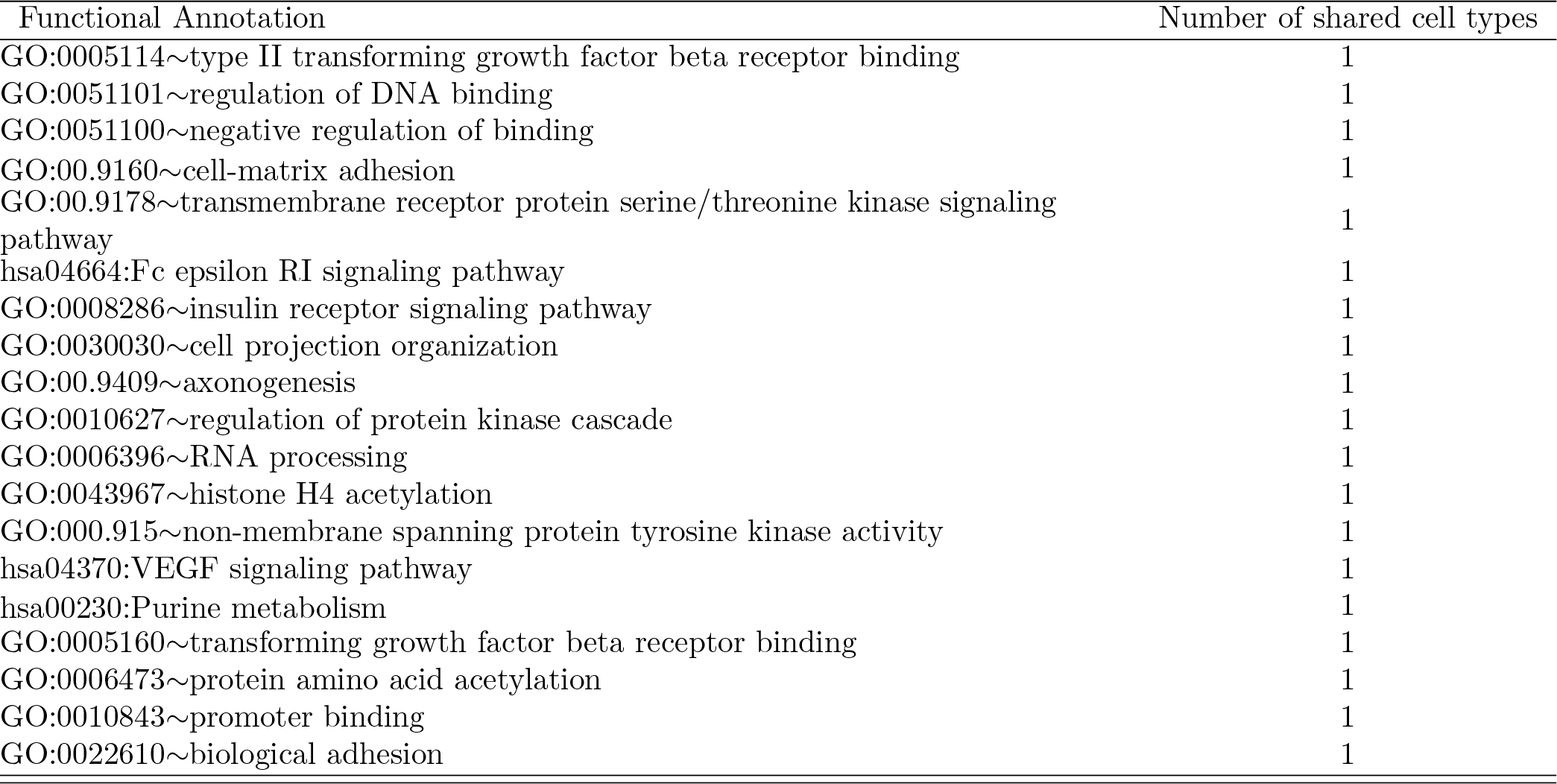

